# Sex Differences in Morphine Sensitivity of Neuroligin-3 Knockout Mice

**DOI:** 10.1101/2024.06.01.596965

**Authors:** Dieter D. Brandner, Mohammed A. Mashal, Nicola M. Grissom, Patrick E. Rothwell

## Abstract

Sex has a strong influence on the prevalence and course of brain conditions, including autism spectrum disorders. The mechanistic basis for these sex differences remains poorly understood, due in part to historical bias in biomedical research favoring analysis of male subjects, and the exclusion of female subjects. For example, studies of male mice carrying autism-associated mutations in neuroligin-3 are over-represented in the literature, including our own prior work showing diminished responses to chronic morphine exposure in male neuroligin-3 knockout mice. We therefore studied how constitutive and conditional genetic knockout of neuroligin-3 affects morphine sensitivity of female mice. In contrast to male mice, female neuroligin-3 knockout mice showed normal psychomotor sensitization after chronic morphine exposure. However, in the absence of neuroligin-3 expression, both female and male mice show a similar change in the topography of locomotor stimulation produced by morphine. Conditional genetic deletion of neuroligin-3 from dopamine neurons increased the locomotor response of female mice to high doses of morphine, contrasting with the decrease in psychomotor sensitization caused by the same manipulation in male mice. Together, our data reveal that knockout of neuroligin-3 has both common and distinct effects on morphine sensitivity in female and male mice. These results also support the notion that female sex can confer resilience against the impact of autism-associated gene variants.

## INTRODUCTION

Sex-related mechanisms, including genetic and hormonal factors, have important influences on cognitive function and diversity (Grissom et al. 2024). These same mechanisms likely contribute to sex differences in the prevalence and course of many brain conditions, including autism spectrum disorders (Werling and Geschwind 2013). Autism spectrum disorders are diagnosed much more frequently in males, due in part to diagnostic gender bias (Loomes et al. 2017). However, sex-related biological mechanisms also appear to protect females from autism-associated genetic variants (Cosgrove et al. 2007). In studies using rodents as an experimental system, a given autism-associated genetic variant can produce male-specific deficits in behavior and brain function (Grissom et al. 2018; Kight et al. 2021). However, a deeper understanding of sex-related influences on the etiology of autism spectrum disorders has been constrained by a large number of studies focused solely on male subjects. This is emblematic of a larger bias favoring male over female research subjects that has pervaded neuroscience research for many years (Mamlouk et al. 2020), and is particularly problematic in studies of autism spectrum disorders and other brain conditions with a strong sex bias.

A clear illustration of this problem comes from autism-associated genetic variants in neuroligin-3 (NL3), a postsynaptic cell adhesion molecule that sculpts synaptic structure and function through interactions with presynaptic binding partners (Uchigashima et al. 2021). NL3 gene variants associated with autism spectrum disorders include both point mutations like the R451C substitution (Jamain et al. 2003), as well as exonic deletions (Levy et al. 2011; Sanders et al. 2011). In studies of mice carrying these genetic variants, some phenotypes reported in NL3^R451C/y^ mutant mice have not been observed in NL3^-/y^ constitutive knockout mice. These phenotypes may represent a gain-of-function related to the R451C point mutation, and include reduced social interaction as well as enhanced spatial learning, with variable expression depending on genetic background (Cao et al. 2018; Chadman et al. 2008; Etherton et al. 2011; Jaramillo et al. 2018; Jaramillo et al. 2014; Tabuchi et al. 2007). Conversely, common phenotypes observed in both NL3^R451C/y^ and NL3^-/y^ mutant mice are likely due to loss of NL3 function. These include locomotor hyperactivity in the open field test and enhanced motor learning on the accelerating rotarod, which manifest on multiple genetic backgrounds (Rothwell et al. 2014).

Additional loss-of-function effects described in NL3^-/y^ knockout mice (but not assessed in NL3^R451C/y^ mutant mice) include decreased psychomotor sensitization after chronic morphine exposure (Brandner et al. 2023), and less rewarding properties of social interaction using conditioned place preference (Bariselli et al. 2018). Both of these effects appear to specifically involve NL3 expression by dopamine neurons, where loss of NL3 impairs oxytocin signaling (Hornberg et al. 2020). NL3 knockout mice can also modify the social behavior of wildtype littermates (Kalbassi et al. 2017). However, with few exceptions (e.g., Chadman et al. 2008; Kalbassi et al. 2017), nearly all of these studies have characterized only male mice carrying NL3 mutations. This places an inherent limitation on our understanding of NL3 function, as it remains unclear if and how these NL3 mutations impact female subjects.

For X-linked genes like NL3, male wild-type and hemizygous knockout littermates can be generated from a single breeding scheme. However, the generation female wild-type and homozygous knockout mice requires distinct breeding schemes that each yield heterozygous littermates (e.g., Chadman et al. 2008; Kalbassi et al. 2017). Using these distinct breeding schemes (Figure 1), we have characterized the behavioral sensitivity of female NL3 knockout mice to acute and chronic morphine exposure, to complement our prior study of male NL3 knockout mice (Brandner et al. 2023). We find that female and male NL3 knockout mice exhibit some similarity but also important differences in terms of morphine sensitivity, suggesting sex-related molecular mechanisms for opioid-evoked plasticity involving synaptic cell adhesion molecules. By demonstrating sex-related influences of NL3 knockout on opioid sensitivity, these data highlight a new dimension of male vulnerability and female resilience to genetic variation associated with neuropsychiatric conditions.

**Figure 1.**
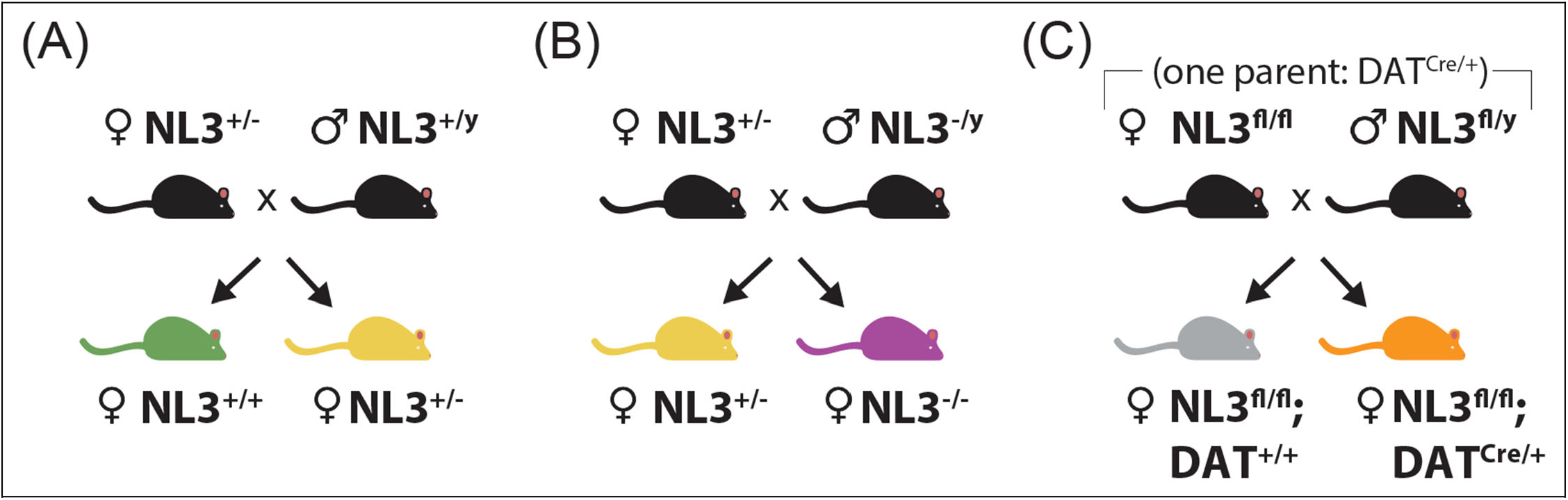
Breeding strategies used to generate female mice with neuroligin-3 (NL3) mutations for behavioral analysis. (A) NL3^+/-^ dams crossed with NL3^+/y^ sires, generating female littermates that are either NL3^+/-^ or NL3^+/+^. (B) NL3^+/-^ dams crossed with NL3^-/y^ sires, generating female littermates that are either NL3^+/-^ or NL3^-/-^. (C) NL3^fl/fl^ dams crossed with NL3^fl/y^ sires, with one parent carrying a single copy of the DAT-IRES-Cre allele (DAT^Cre/+^), generating female NL3^fl/fl^ littermates that either carry a single copy of Cre (DAT^Cre/+^) or lack Cre expression (DAT^+/+^).

## METHODS

### Subjects

Mice were housed in groups of two to five per cage, on a 12-hour light cycle (0600–1800 h) at ∼23°C with ad libitum access to food and water. Experimental procedures were conducted between 1000h-1600h, were approved by to the University of Minnesota Institutional Animal Care and Use Committee, and observed the NIH *Guidelines for the Care and Use of Laboratory Animals*. All mouse lines were maintained on a C57Bl/6J genetic background. The generation of constitutive NL3 knockout animals (The Jackson Laboratory Strain #008394) has been previously described (Tabuchi et al. 2007). Female NL3 knockout mice were generated using one of two different breeding schemes. In the first scheme, NL3^+/y^ sires were crossed to NL3^+/-^ dams, generating female offspring that were either NL3^+/-^ or NL3^+/+^ (Figure 1A). In the second scheme, NL3^-/y^ sires were crossed to NL3^+/-^ dams, generating female offspring that were either NL3^+/-^ or NL3^-/-^ (Figure 1B). The generation of conditional NL3 knockout animals has been previously described (Rothwell et al. 2014). To conditionally delete NL3 expression in dopamine neurons, this line was crossed with DAT-IRES-Cre knock-in (Backman et al. 2006) and then maintained by breeding NL3^fl/y^ sires and NL3^fl/fl^ dams, with one of the two parents carrying a single copy of DAT-IRES-Cre (Figure 1C). This generated female NL3^fl/fl^ offspring that carried a single copy of Cre (DAT^Cre/+^), along with littermate control mice lacking Cre expression (DAT^+/+^).

### Morphine administration

Morphine hydrochloride (Mallinckrodt) was dissolved in sterile 0.9% saline and delivered subcutaneously (5 mL/kg) by bolus injection. The range of morphine doses (2, 6.32, or 20 mg/kg) has previously been used by our group to measure the acute psychomotor response to morphine, as well as long-lasting psychomotor sensitization (Brandner et al. 2023; Lefevre et al. 2020; Toddes et al. 2021). The day prior to the first acute morphine injection, mice were habituated to the behavior apparatus and given an equivalent volume of 0.9% saline subcutaneously. This was followed by exposure to increasing morphine doses of 2, 6.32, and 20 mg/kg morphine on consecutive days (Figure 2; “Acute”). Psychomotor sensitization consisted of seven consecutive doses of daily morphine administered subcutaneously at 20 mg/kg. To minimize the contextual effects of the behavior apparatus on psychomotor sensitization, locomotion was only measured after the first and last injection of 20 mg/kg morphine on Days 1 and 7 (Figure 2; “Sensitization”). Three weeks after the end of morphine sensitization, we delivered morphine challenge injections in a manner analogous to acute dose–response (Figure 2; “Challenge”).

**Figure 2.**
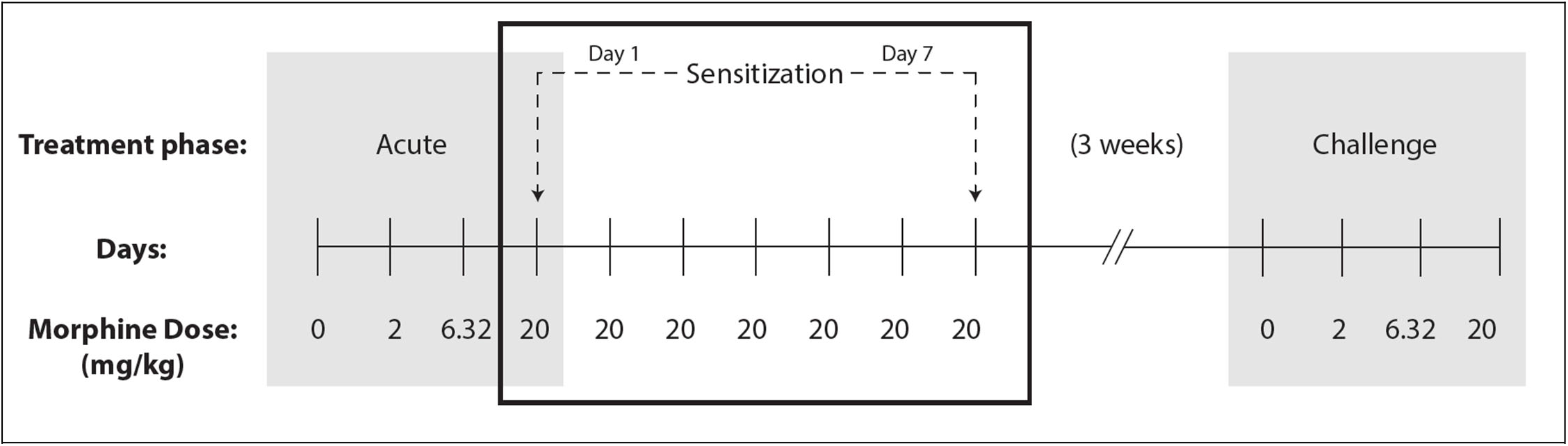
Experimental timeline for morphine administration. Acute exposure began with injection of 0, 2, 6.32, and 20 mg/kg morphine on consecutive days. Sensitization continued with seven daily injections of 20 mg/kg morphine, with measurement of locomotor response (dashed lines) after the first injection (Day 1) and last injection (Day 7). Three weeks after the end of sensitization, morphine challenge was conducted using ascending doses in a manner analogous to acute dose-response.

### Behavioral Analysis

As previously described (Brandner et al. 2023; Lefevre et al. 2020; Toddes et al. 2021), we tested open field locomotor activity in a clear plexiglass arena (ENV510, Med Associates) within a sound-attenuating chamber. Locomotor test sessions began immediately after drug injection and lasted 60 minutes. The center zone of the open field arena was defined based on the central 75% in each dimension, and distance travelled both within and outside this center zone was automatically calculated using Activity Monitor software (Med Associates).

### Statistical Analysis

Sample sizes are indicated in figure legends and graphs that display individual data points, along with measures of central tendency (mean) and variability (standard error of the mean). Data were analyzed in factorial ANOVA models using GraphPad Prism 10, with repeated measures on within-subject factors or mixed-effect models to handle missing data. For main effects or interactions involving repeated measures, the Greenhouse-Geisser correction was applied to control for potential violations of the sphericity assumption. This correction reduces the degrees of freedom, resulting in non-integer values. Significant interactions were decomposed by analyzing simple effects (i.e., the effect of one variable at each level of the other variable). Significant main effects were analyzed using LSD post-hoc tests. The Type I error rate was set to α = 0.05 (two-tailed) for all comparisons.

## RESULTS

### Effects of Heterozygous NL3 Deletion in Female Constitutive Knockout Mice

In a prior study, we found that male mice with constitutive genetic deletion of NL3 had normal acute responses to morphine, but a persistent reduction of psychomotor sensitization after daily morphine injections (Brandner et al. 2023). To study behavioral responses to morphine in female mice with NL3 mutations, we first crossed NL3^+/y^ wild-type males with NL3^+/-^ heterozygous knockout females (Figure 1A). Offspring from this cross included female NL3^+/-^ heterozygous knockouts, and a littermate control group of female NL3^+/+^ wild-types, which were compared in behavioral studies.

We first measured acute locomotor responses to ascending doses of morphine (0-20 mg/kg), followed by daily exposure to a high dose (20 mg/kg), and concluded with a dose-response “challenge” following three weeks of withdrawal (Figure 2). After acute morphine administration (Figure 3A), all mice exhibited a dose-dependent increase in locomotor activity, as indicated by a significant main effect of Dose (F_1.46,27.82_ = 319.8, *P*<0.0001). However, there was no main effect of Genotype (F_1,19_ < 1), and no Dose x Genotype interaction (F_3,57_ = 2.46, *p* = 0.072). This indicates that female NL3^+/+^ and NL3^+/-^ mice had a similar locomotor response to acute morphine administration.

**Figure 3.**
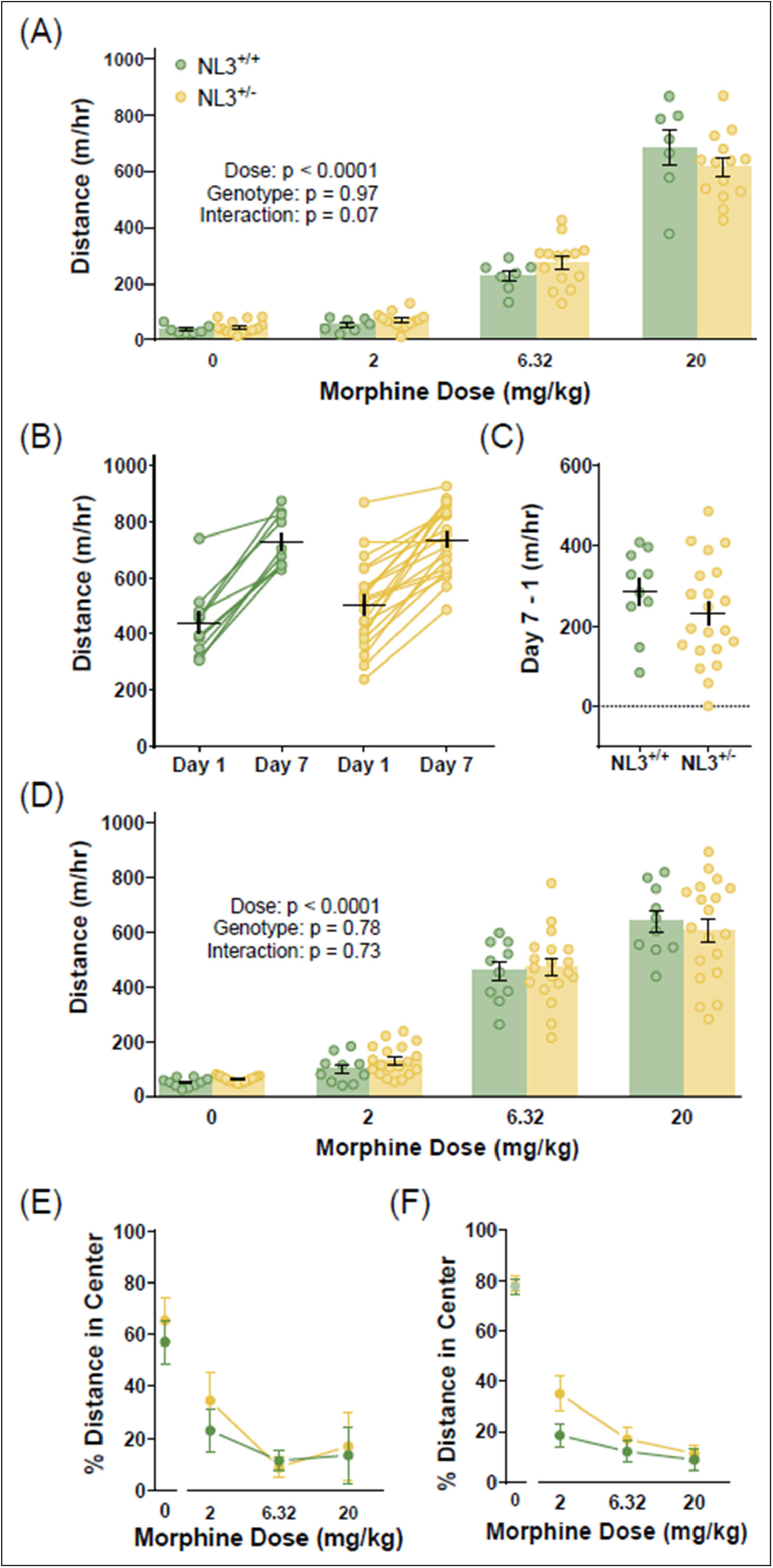
Responses to acute and chronic morphine in female NL3 heterozygous knockout mice and their wild-type littermates. (A) Acute locomotor response to injection of ascending morphine doses. (B) Locomotor response to 20 mg/kg morphine after the first and last daily injection. (C) Change in locomotor response to 20 mg/kg morphine following daily injection. (D) Locomotor response to morphine challenge conducted three weeks after the end of sensitization. (E, F) Percentage of total distance travelled in the central area after acute (E) or chronic (F) morphine exposure. Sample size: n = 10 (NL3^+/+^) and 20 (NL3^+/-^).

After seven daily injections of 20 mg/kg morphine (Figure 3B), all mice exhibited locomotor sensitization, as indicated by a significant main effect of Day (F_1,29_ = 121.8, *p* < 0.0001). However, there was no main effect of Genotype (F_1,29_ < 1), and no Day x Genotype interaction (F_1,29_ = 1.427, *p* = 0.24). This indicates that female NL3^+/+^ and NL3^+/-^ mice developed a similar degree of psychomotor sensitization, which was also apparent after computing the change in distance travelled between Days 1 and 7 for each individual animal (Figure 3C; t_29_ = 1.19, *p* = 0.24). The durability of psychomotor sensitization was also similar 21 days later, when the same cohort of animal were challenged with increasing doses of morphine (Figure 3D). There was significant main effect of Dose (F_1.79,48.32_ = 183.8, *p* < 0.0001), but no main effect of Genotype (F_1, 27_ < 1), and no Dose x Genotype interaction (F_3,81_ < 1).

After morphine injection, the locomotor hyperactivity exhibited by wild-type mice normally leads to a path of travel along the perimeter of the activity chamber. We previously quantified the topography of locomotor activity by calculating the percentage of total distance traveled through the interior of the chamber (Brandner et al. 2023), and repeated this analysis using the current data from female NL3^+/+^ and NL3^+/-^ mice. For acute morphine (Figure 3E), there was no baseline difference between groups after saline injection (t_11_ < 1), and no significant main effects or interactions after acute morphine exposure. For morphine challenge (Figure 3F), there was no baseline difference between groups after saline injection (t_21_ < 1). Morphine challenge caused a dose-dependent decrease in the percent of distance travelled through the center of the chamber (F_1.85,38.86_ = 8.697, *p* = 0.001), but there was no main effect of Genotype (F_1,42_ = 2.21, *p* = 0.15), and no Dose x Genotype interaction (F_2,42_ = 1.607, *p* = 0.21). Heterozygous constitutive knockout of NL3 thus had no significant impact on behavioral responses to morphine in female mice.

### Effects of Homozygous NL3 Deletion in Female Constitutive Knockout Mice

To determine if homozygous constitutive knockout of NL3 affected behavioral sensitivity to morphine in female mice, we crossed NL3^-/y^ male knockouts with NL3^+/-^ heterozygous knockout females (Figure 1B). Offspring from this cross included female NL3^-/-^ homozygous knockouts, and littermate NL3^+/-^ females that served as a control group for comparison, since the behavioral sensitivity of NL3^+/-^ heterozygous knockout females was similar to NL3^+/+^ wild-type females (Figure 3).

After acute morphine administration (Figure 4A), all mice exhibited a dose-dependent increase in locomotor activity, as indicated by a significant main effect of Dose (F_1.69, 30.39_ = 270.9, *p* < 0.0001). However, there was no main effect of Genotype (F_1, 18_ = 1.08, *p* = 0.31), and no Dose x Genotype interaction (F_3, 54_ = 2.51, *p* = 0.069). This indicates that female NL3^+/-^ and NL3^-/-^ mice had a similar locomotor response to acute morphine administration.

**Figure 4.**
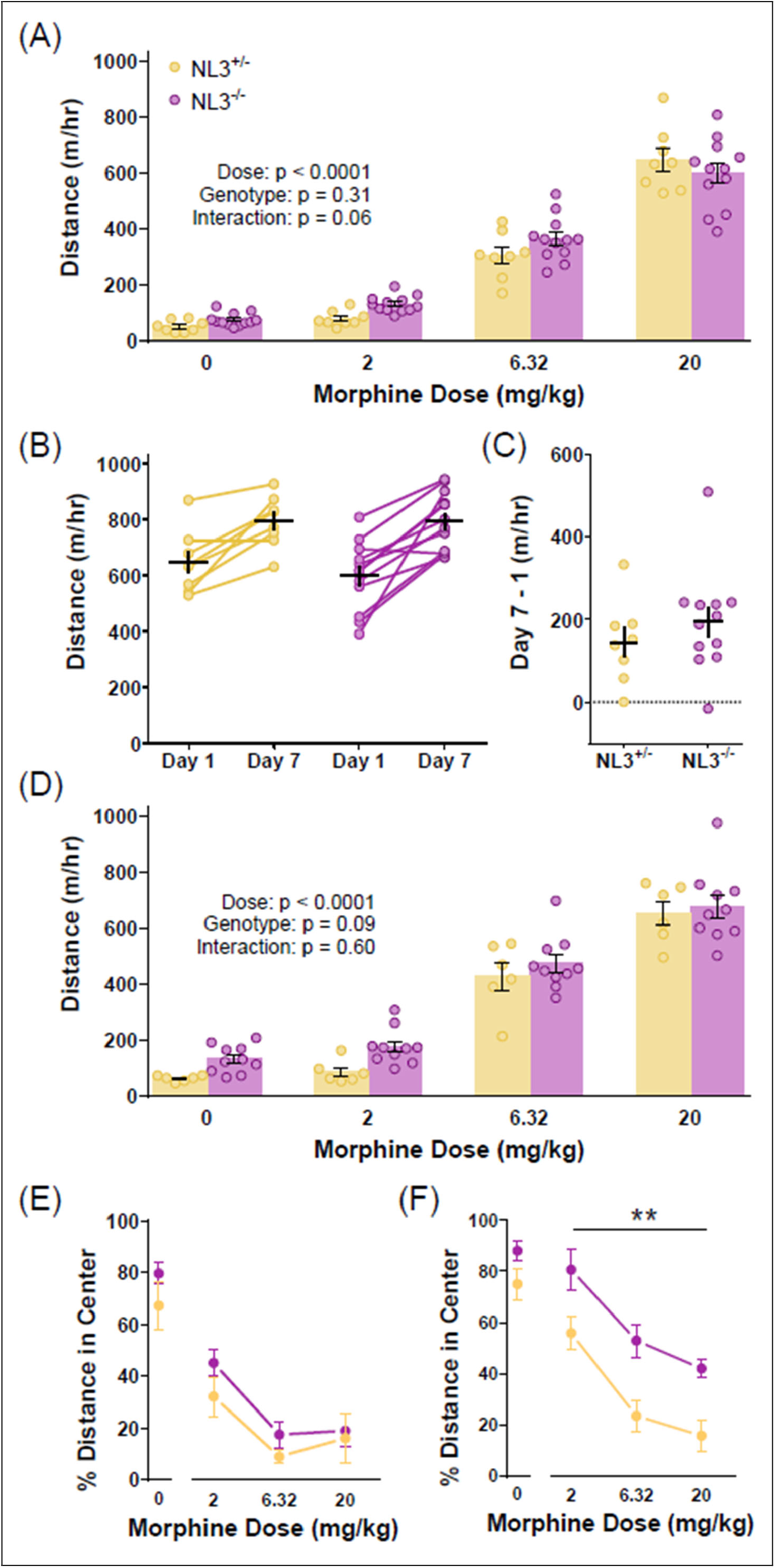
Responses to acute and chronic morphine in female NL3 homozygous knockout mice and their heterozygous knockout littermates. (A) Acute locomotor response to injection of ascending morphine doses. (B) Locomotor response to 20 mg/kg morphine after the first and last daily injection. (C) Change in locomotor response to 20 mg/kg morphine following daily injection. (D) Locomotor response to morphine challenge conducted three weeks after the end of sensitization. (E, F) Percentage of total distance travelled in the central area after acute (E) or chronic (F) morphine exposure. Sample size: n = 8 (NL3^+/-^

After seven daily injections of 20 mg/kg morphine (Figure 4B), all mice exhibited locomotor sensitization, as indicated by a significant main effect of Day (F_1, 18_ = 41.04, *p* < 0.0001). However, there was no main effect of Genotype (F_1, 18_ < 1), and no Day x Genotype interaction (F_1, 18_ < 1). This indicates that female NL3^+/-^ and NL3^-/-^ mice developed a similar degree of psychomotor sensitization, which was also apparent after computing the change in distance travelled between Days 1 and 7 for each individual animal (Figure 4C; t_18_ < 1). The durability of psychomotor sensitization was also similar upon morphine challenge 21 days later (Figure 4D). There was significant main effect of Dose (F_1.94, 27.21_ = 214.1, *p* < 0.0001), but no main effect of Genotype (F_1, 14_ = 3.12, *p* =0.099), and no Dose x Genotype interaction (F_3, 42_ < 1). The intact psychomotor sensitization in female NL3^-/-^ knockout mice contrasts with the attenuated psychomotor sensitization we previously reported in male NL3^-/y^ knockout mice (Brandner et al. 2023).

Male NL3^-/y^ knockout mice also exhibit a more curvilinear path of travel after morphine injection, bypassing the corners of the chamber and instead traveling through more central areas (Brandner et al. 2023). We thus quantified the percentage of total distance traveled through the interior of the chamber using the current data for female NL3^+/-^ and NL3^-/-^ mice. For acute morphine (Figure 4E), there was no baseline difference between groups after saline injection (t_14_ = 1.34, *p* = 0.20). Acute morphine caused a dose-dependent decrease in the percent of distance travelled through the center of the chamber (F_1.42,20.60_ = 12.65, *p* = 0.0007), but there was no main effect of Genotype (F_1,14_ = 1.66, *p* = 0.22) and no Dose x Genotype interaction (F_2,28_ < 1). For morphine challenge (Figure 4F), there was no baseline difference between groups after saline injection (t_14_ = 1.89, *p* = 0.079). Morphine challenge caused a dose-dependent decrease in the percent of distance travelled through the center of the chamber (F_1.63,22.78_ = 40.3, *p* < 0.0001). There was also a significant main effect of Genotype (F_1,14_= 13.99, *p* = 0.0022), indicating that female NL3^-/-^ knockout mice travelled more through the center regardless of morphine dose, as there was no Dose x Genotype interaction (F_2,28_ < 1). This change in locomotor topography after morphine challenge is thus shared in common between female NL3^-/-^ knockout mice and male NL3^-/y^ knockout mice (Brandner et al. 2023).

### Effects of Conditional NL3 Deletion from Dopamine Neurons in Female Mice

In our prior study of male mice, we found that conditional genetic deletion of NL3 from dopamine neurons was sufficient to attenuate psychomotor sensitization to morphine (Brandner et al. 2023). To repeat these behavioral experiments in female mice, we used conditional NL3 knockout mice with loxP sites flanking exons 2 and 3 of the *Nlgn3* gene (Rothwell et al. 2014). We crossed NL3^fl/fl^ females and NL3^fl/y^ males, with one parent also carrying a single copy of DAT-IRES-Cre (DAT^Cre/+^). This breeding strategy generated female offspring that were all NL3^fl/fl^ (Figure 1C), with 50% carrying DAT-Cre (NL3^fl/fl^;DAT^Cre/+^ experimental group) and 50% lacking DAT-Cre (NL3^fl/fl^;DAT^+/+^ littermate control group).

After acute morphine administration (Figure 5A), all mice exhibited a dose-dependent increase in locomotor activity, as indicated by a significant main effect of Dose (F_1.87,48.66_ = 124.1, *p* < 0.0001). There was also a significant main effect of Genotype (F_1,26_ = 6.438, *p* = 0.0175), indicating a greater distance traveled in mutant mice regardless of dose, as there was no Dose x Genotype interaction (F_3,78_ = 2.23, *p* = 0.091). This main effect of Genotype for acute morphine exposure was interesting, as it was not previously observed in either female NL3 mutants (Figures 3-4) or male NL3 mutants (Brandner et al. 2023).

**Figure 5.**
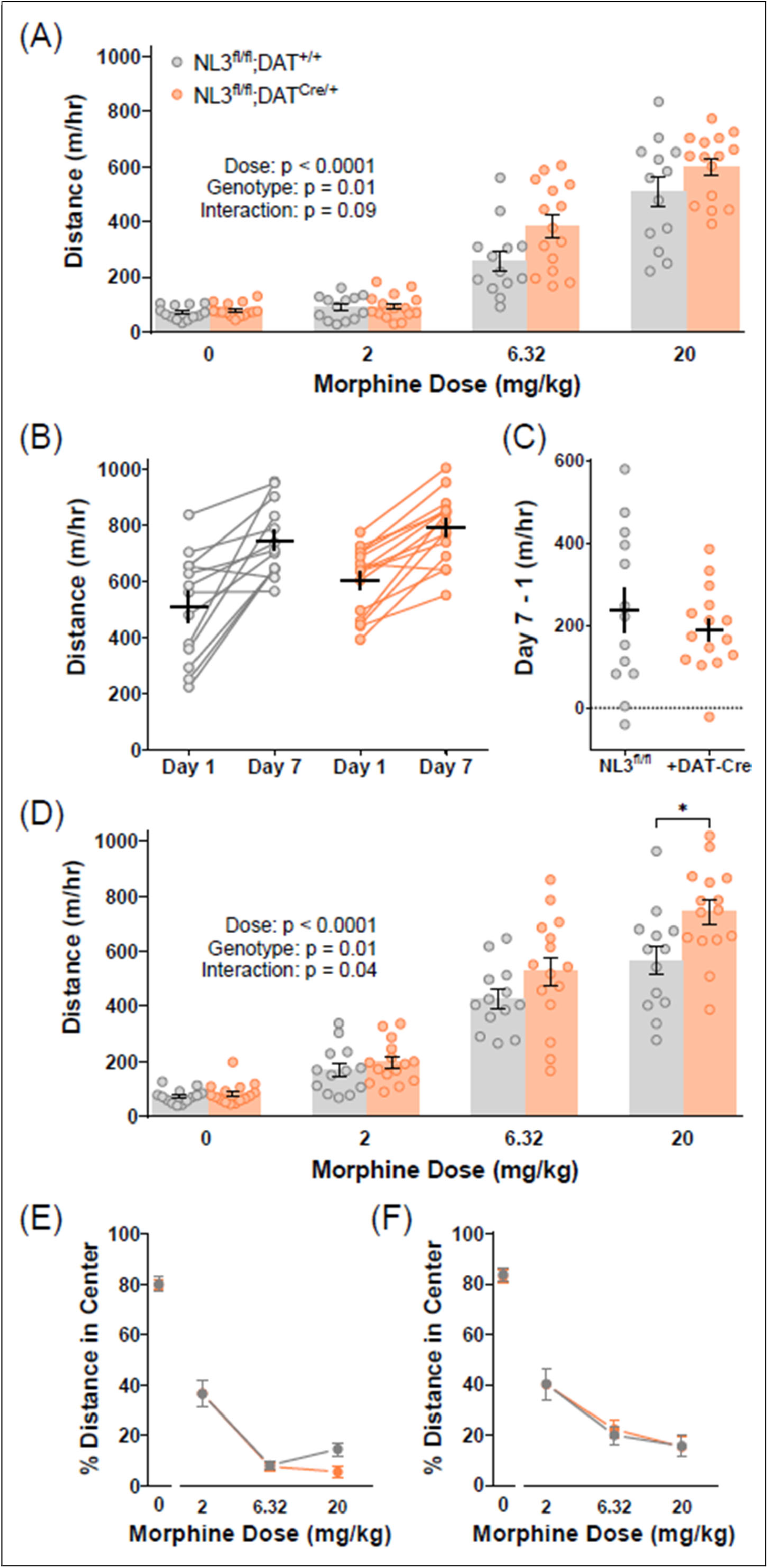
Responses to acute and chronic morphine in female mice after conditional NL3 deletion from dopamine neurons. (A) Acute locomotor response to injection of ascending morphine doses. (B) Locomotor response to 20 mg/kg morphine after the first and last daily injection. (C) Change in locomotor response to 20 mg/kg morphine following daily injection. (D) Locomotor response to morphine challenge conducted three weeks after the end of sensitization. (E, F) Percentage of total distance travelled in the central area after acute (E) or chronic (F) morphine exposure. Sample size: n = 13 (NL3^fl/fl^;DAT^+/+^) and 15 (NL3^fl/fl^;DAT^Cre/+^).

After seven daily injections of 20 mg/kg morphine (Figure 5B), all mice exhibited locomotor sensitization, as indicated by a significant main effect of Day (F_1,26_ = 55.61, *p* < 0.0001). However, there was no main effect of Genotype (F_1,26_ = 2.23, *p* = 0.15), and no Day x Genotype interaction (F_1, 26_ < 1). This indicates that female NL3^fl/fl^ mice developed a similar degree of psychomotor sensitization regardless of DAT-Cre expression, which was also apparent after computing the change in distance travelled between Days 1 and 7 for each individual animal (Figure 5C; t_26_ < 1). Following morphine challenge (Figure 5D), there were significant main effects of Dose (F_1.95,50.70_ = 137.1, *p* < 0.0001) and Genotype (F_1,26_ = 6.65, *p* = 0.016), as well as a significant Dose x Genotype interaction (F_3,78_ = 2.83, *p* = 0.044). This interaction indicated the simple effect of Genotype was most robust after challenge with 20 mg/kg morphine (t_24_ = 2.59, *p* = 0.016), with greater distance traveled in mice with DAT-Cre expression, though a similar pattern was present after challenge with 6.32 mg/kg morphine as well as acute exposure to 6.32 and 20 mg/kg morphine (Figure 5A). Female NL3^fl/fl^;DAT^Cre/+^ mice thus appear to have an enhanced locomotor response to higher doses of morphine regardless of chronic treatment, which is distinct from the phenotype of male NL3^fl/y^;DAT^Cre/+^ mice (Brandner et al. 2023).

In contrast, the path of travel following morphine exposure in female NL3^fl/fl^ mice was not affected by DAT-Cre expression. For acute morphine (Figure 5E), there was no baseline difference between groups after saline injection (t_26_ < 1). Acute morphine caused a dose-dependent decrease in the percent of distance travelled through the center of the chamber (F_1.30,33.92_ = 53.98, *p* < 0.0001), but there was no main effect of Genotype (F_1,26_ < 1) and no Dose x Genotype interaction (F_2,52_ = 1.38, *p* = 0.26). For morphine challenge (Figure 5F), there was no baseline difference between groups after saline injection (t_26_ < 1). Morphine challenge caused a dose-dependent decrease in the percent of distance travelled through the center of the chamber (F_1.64,42.77_ = 23.18, *p* < 0.0001), but there was no main effect of Genotype (F_1,26_ < 1), and no Dose x Genotype interaction (F_2,52_ < 1). Combined with our prior analysis of male mice (Brandner et al. 2023), these results show that conditional deletion of NL3 from dopamine neurons thus has no effect on locomotor topography following morphine injection in mice of either sex.

## DISCUSSION

Exogenous opioid exposure has profound effects on brain function and behavior, causing adaptations in dopaminergic circuits that include changes in synaptic structure and function. Synaptic cell adhesion molecules like NL3 are prime candidates for mediating opioid-evoked plasticity. In male mice, we previously showed that NL3 in dopaminergic circuits promotes behavioral and neurobiological adaptations to chronic morphine exposure (Brandner et al. 2023). However, female mice have not been included in many prior studies of NL3 function, including our own. Our current data highlight intriguing sex differences in morphine sensitivity of NL3 knockout mice (summarized in Table 1), which add a growing body of literature related to sex differences in neural mechanisms mediating substance use disorders (Becker and Chartoff 2019).

**Table 1.**
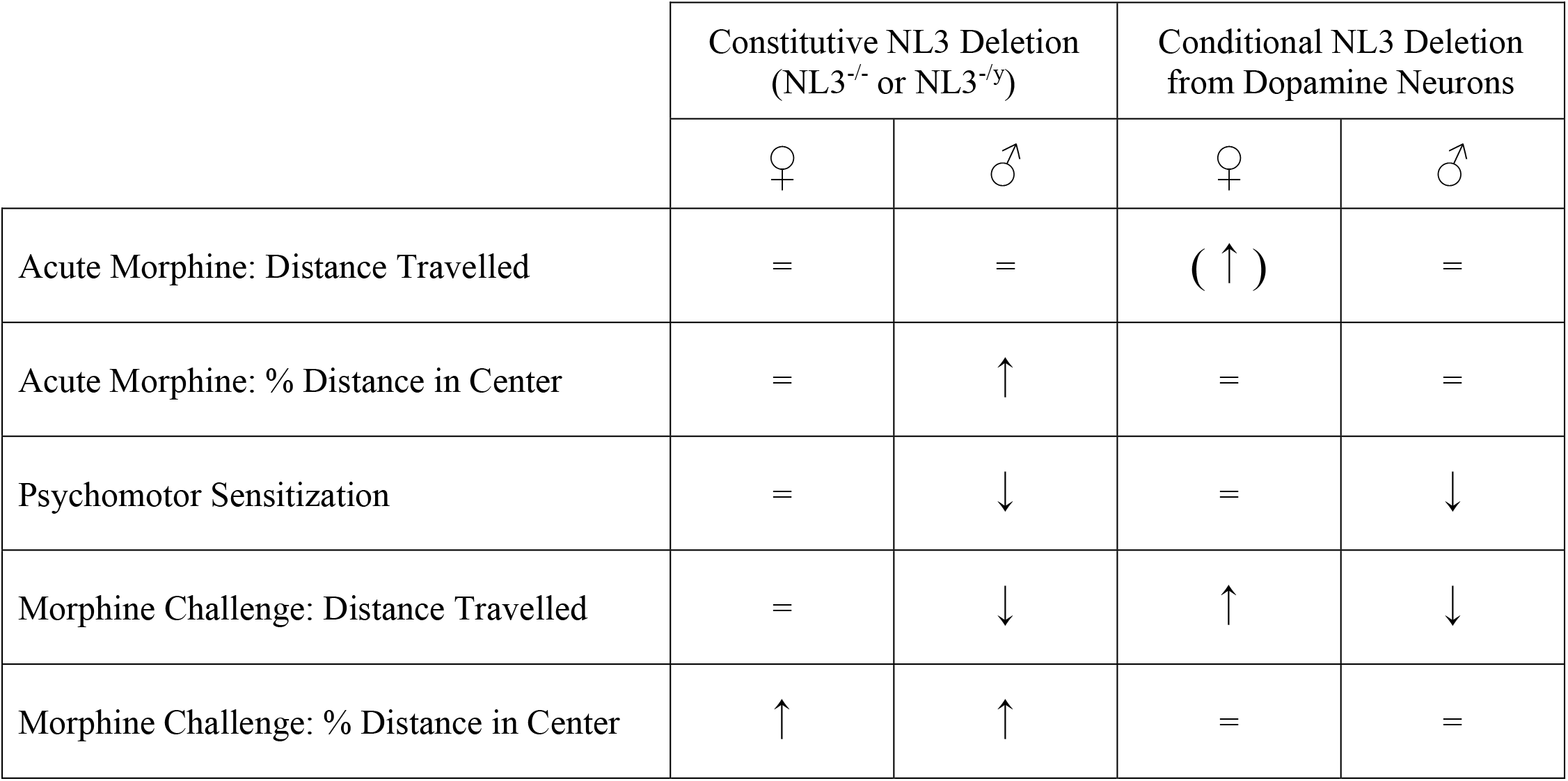
Changes in different measures of morphine sensitivity in female mice (this study) and male mice (Brandner et al. 2023), comparing NL3 mutants to littermate controls.

In male mice, constitutive genetic knockout of NL3 caused a persistent reduction of psychomotor sensitization after daily morphine injections, which was recapitulated by conditional genetic deletion of NL3 from dopamine neurons. In female mice, however, constitutive genetic knockout of NL3 did not impact the development of psychomotor sensitization after daily morphine injection. This was true for both heterozygous NL3^+/-^ knockouts compared to wildtype littermates (Figure 3), and homozygous NL3^-/-^ knockouts compared to heterozygous NL3^+/-^ littermates (Figure 4). Female mice were thus “protected” against this phenotype of NL3 deletion, similar to enhanced motor learning on the accelerating rotarod, a phenotype seen in male NL3^R451C/y^ and NL3^-/y^ mutant mice but absent in female mice carrying the R451C mutation in NL3 (Chadman et al. 2008; Rothwell et al. 2014).

However, genetic deletion of NL3 did impact other behavioral responses to morphine in female mice. Homozygous NL3^-/-^ knockout females exhibited a change in locomotor topography after challenge injections of morphine, traveling through more central areas of the chamber rather than moving along the perimeter of the chamber (Figure 4F). Male NL3 knockout mice exhibit a similar and robust change in locomotor topography after challenge injections in morphine (Brandner et al. 2023), though male NL3 knockout mice also showed a change in locomotor topography after acute morphine exposure, which was less apparent in female NL3 knockout mice (Figure 4E). In our prior study of male mice, conditional deletion of NL3 from cells that express the Drd1 dopamine receptor reproduced the change in locomotor topography after challenge injections of morphine, even though this manipulation had no effect on psychomotor sensitization (Brandner et al. 2023). In future studies, it will be interesting to determine how female mice respond following conditional deletion of NL3 from cells that express the Drd1 dopamine receptor, including evaluation of locomotor topography after morphine challenge as well as other behavioral phenotypes produced by this genetic manipulation in male mice (Rothwell et al. 2014).

In this study, we further evaluated female mice with conditional deletion of NL3 from dopamine neurons, by crossing a conditional NL3 knockout line with DAT-Cre. Previous studies of male mice have described regulation of social behavior by NL3 expression in dopamine neurons (Bariselli et al. 2018; Hornberg et al. 2020), and we previously observed that conditional deletion of NL3 from dopamine neurons phenocopied the decrease in morphine-evoked psychomotor sensitization seen after constitutive NL3 deletion in samples of male mice (Brandner et al. 2023). However, female mice showed a unique behavioral profile following conditional deletion of NL3 from dopamine neurons, with an enhanced locomotor response to higher doses of morphine that appeared to be present regardless of chronic morphine treatment. This particular genetic manipulation thus produced distinct and somewhat opposite effects on morphine sensitivity in mice of each sex, highlighting the potential for great complexity in the molecular code of synaptic cell adhesion that regulates behavioral responses to opioids and other misused drugs. It will be interesting for future studies to determine if NL3 expression in dopamine neurons also regulates social behavior and reward in a sex-dependent fashion, as this could have important therapeutic implications for treatments targeting cellular dysfunction in dopamine neurons after loss of NL3 (Hornberg et al. 2020).

The mechanistic basis for sex differences in the impact of neuroligin-3 deletion also merits further study. One possibility is that sex-related mechanisms influence the ability of other members of the neuroligin family to compensate for the constitutive neuroligin-3 knockout. Neuroligin-3 is localized to both excitatory and inhibitory synapses (Budreck and Scheiffele 2007), so upregulation of neuroligin-1 and neuroligin-2 could be a source of compensation at excitatory synapses and inhibitory synapses, respectively (Song et al. 1999; Varoqueaux et al. 2004). While there is no evidence for compensatory upregulation of other neuroligins in forebrain tissue of male constitutive neuroligin-3 knockout mice (Tabuchi et al. 2007), this analysis has not been conducted in females and results could differ. Another possibility is that the pattern and level of neuroligin-3 expression across different brain regions and cell types is influenced by sex-related mechanisms. However, our data show that a specific molecular manipulation of neuroligin-3 (deletion from dopamine neurons) can have seemingly opposite effects on morphine sensitivity. This result hints at the possibility of more fundamental sex differences in the cell biology of neuroligin-3 function within a given cell type, which could relate to sex-dependent interactions with various intracellular and extracellular binding partners (e.g., Yoshida et al. 2021).

In closing, we note that new therapeutic opportunities may arise from careful attention to sex as a biological variable in the context of genetic variants associated with brain conditions. A major concern about bias towards research on male subjects is that it can drive the development of therapeutic strategies that are less effective, ineffective, or even have adverse side effects in female subjects (Dalla et al. 2024; Shansky and Murphy 2021). Our results suggest an additional opportunity that may be missed by excluding female subjects from research on autism-associated genetic variants: that a clearer biological understanding of female resilience could inspire new therapeutic strategies that promote resilience in male subjects.

## Acknowledgements

This research was supported by the University of Minnesota’s MnDRIVE (Minnesota’s Discovery, Research and Innovation Economy) initiative, as well as National Institutes of Health grants T32 DA052109 (DDB), F30 DA007234 (DDB), R01 MH123661 (NMG), R01 DA048946 (PER), and P50 MH119569 (NMG/PER). The authors have no conflicts of interest to declare.

## REFERENCES

Backman CM, Malik N, Zhang Y, Shan L, Grinberg A, Hoffer BJ, Westphal H, Tomac AC (2006) Characterization of a mouse strain expressing Cre recombinase from the 3’ untranslated region of the dopamine transporter locus. Genesis 44: 383–90.

Bariselli S, Hornberg H, Prevost-Solie C, Musardo S, Hatstatt-Burkle L, Scheiffele P, Bellone C (2018) Role of VTA dopamine neurons and neuroligin 3 in sociability traits related to nonfamiliar conspecific interaction. Nat Commun 9: 3173.

Becker JB, Chartoff E (2019) Sex differences in neural mechanisms mediating reward and addiction. Neuropsychopharmacology 44: 166–183.

Brandner DD, Retzlaff CL, Kocharian A, Stieve BJ, Mashal MA, Mermelstein PG, Rothwell PE (2023) Neuroligin-3 in dopaminergic circuits promotes behavioural and neurobiological adaptations to chronic morphine exposure. Addict Biol 28: e13247.

Budreck EC, Scheiffele P (2007) Neuroligin-3 is a neuronal adhesion protein at GABAergic and glutamatergic synapses. Eur J Neurosci 26: 1738–48.

Cao W, Lin S, Xia QQ, D. YL, Yang Q, Zhang MY, Lu YQ, Xu J, Duan SM, Xia J, Feng G, Xu J, Luo JH (2018) Gamma Oscillation Dysfunction in mPFC Leads to Social Deficits in Neuroligin 3 R451C Knockin Mice. Neuron 97: 1253–1260 e7.

Chadman KK, Gong S, Scattoni ML, Boltuck SE, Gandhy SU, Heintz N, Crawley JN (2008) Minimal aberrant behavioral phenotypes of neuroligin-3 R451C knockin mice. Autism Res 1: 147–58.

Cosgrove KP, Mazure CM, Staley JK (2007) Evolving knowledge of sex differences in brain structure, function, and chemistry. Biol Psychiatry 62: 847–55.

Dalla C, Jaric I, Pavlidi P, Hodes GE, Kokras N, Bespalov A, Kas MJ, Steckler T, Kabbaj M, Wurbel H, Marrocco J, Tollkuhn J, Shansky R, Bangasser D, Becker JB, McCarthy M, Ferland-Beckham C (2024) Practical solutions for including sex as a biological variable (SABV) in preclinical neuropsychopharmacological research. J Neurosci Methods 401: 110003.

Etherton M, Foldy C, Sharma M, Tabuchi K, Liu X, Shamloo M, Malenka RC, Sudhof TC (2011) Autism-linked neuroligin-3 R451C mutation differentially alters hippocampal and cortical synaptic function. Proc Natl Acad Sci U S A 108: 13764–9.

Grissom NM, Glewwe N, Chen C, Giglio E (2024) Sex mechanisms as nonbinary influences on cognitive diversity. Horm Behav 162: 105544.

Grissom NM, McKee SE, Schoch H, Bowman N, Havekes R, O’Brien WT, Mahrt E, Siegel S, Commons K, Portfors C, Nickl-Jockschat T, Reyes TM, Abel T (2018) Male-specific deficits in natural reward learning in a mouse model of neurodevelopmental disorders. Mol Psychiatry 23: 544–555.

Hornberg H, Perez-Garci E, Schreiner D, Hatstatt-Burkle L, Magara F, Baudouin S, Matter A, Nacro K, Pecho-Vrieseling E, Scheiffele P (2020) Rescue of oxytocin response and social behaviour in a mouse model of autism. Nature 584: 252–256.

Jamain S, Quach H, Betancur C, Rastam M, Colineaux C, Gillberg IC, Soderstrom H, Giros B, Leboyer M, Gillberg C, Bourgeron T, Paris Autism Research International Sibpair S (2003) Mutations of the X-linked genes encoding neuroligins NLGN3 and NLGN4 are associated with autism. Nat Genet 34: 27–9.

Jaramillo TC, Escamilla CO, Liu S, Peca L, Birnbaum SG, Powell CM (2018) Genetic background effects in Neuroligin-3 mutant mice: Minimal behavioral abnormalities on C57 background. Autism Res 11: 234–244.

Jaramillo TC, Liu S, Pettersen A, Birnbaum SG, Powell CM (2014) Autism-related neuroligin-3 mutation alters social behavior and spatial learning. Autism Res 7: 264–72.

Kalbassi S, Bachmann SO, Cross E, Roberton VH, Baudouin SJ (2017) Male and Female Mice Lacking Neuroligin-3 Modify the Behavior of Their Wild-Type Littermates. eNeuro 4.

Kight KE, Argue KJ, Bumgardner JG, Bardhi K, Waddell J, McCarthy MM (2021) Social behavior in prepubertal neurexin 1alpha deficient rats: A model of neurodevelopmental disorders. Behav Neurosci 135: 782–803.

Lefevre EM, Pisansky MT, Toddes C, Baruffaldi F, Pravetoni M, Tian L, Kono TJY, Rothwell PE (2020) Interruption of continuous opioid exposure exacerbates drug-evoked adaptations in the mesolimbic dopamine system. Neuropsychopharmacology 45: 1781–1792.

Levy D, Ronemus M, Yamrom B, Lee YH, Leotta A, Kendall J, Marks S, Lakshmi B, Pai D, Ye K, Buja A, Krieger A, Yoon S, Troge J, Rodgers L, Iossifov I, Wigler M (2011) Rare de novo and transmitted copy-number variation in autistic spectrum disorders. Neuron 70: 886–97.

Loomes R, Hull L, Mandy WPL (2017) What Is the Male-to-Female Ratio in Autism Spectrum Disorder? A Systematic Review and Meta-Analysis. J Am Acad Child Adolesc Psychiatry 56: 466–474.

Mamlouk GM, Dorris DM, Barrett LR, Meitzen J (2020) Sex bias and omission in neuroscience research is influenced by research model and journal, but not reported NIH funding. Front Neuroendocrinol 57: 100835.

Rothwell PE, Fuccillo MV, Maxeiner S, Hayton SJ, Gokce O, Lim BK, Fowler SC, Malenka RC, Südhof TC (2014) Autism-associated neuroligin-3 mutations commonly impair striatal circuits to boost repetitive behaviors. Cell 158: 198–212.

Sanders SJ, Ercan-Sencicek AG, Hus V, Luo R, Murtha MT, Moreno-De-Luca D, Chu SH, Moreau MP, Gupta AR, Thomson SA, Mason CE, Bilguvar K, Celestino-Soper PB, Choi M, Crawford EL, Davis L, Wright NR, Dhodapkar RM, DiCola M, DiLullo NM, Fernandez TV, Fielding-Singh V, Fishman DO, Frahm S, Garagaloyan R, Goh GS, Kammela S, Klei L, Lowe JK, Lund SC, McGrew AD, Meyer KA, Moffat WJ, Murdoch JD, O’Roak BJ, Ober GT, Pottenger RS, Raubeson MJ, Song Y, Wang Q, Yaspan BL, Yu TW, Yurkiewicz IR, Beaudet AL, Cantor RM, Curland M, Grice DE, Gunel M, Lifton RP, Mane SM, Martin DM, Shaw CA, Sheldon M, Tischfield JA, Walsh CA, Morrow EM, Ledbetter DH, Fombonne E, Lord C, Martin CL, Brooks AI, Sutcliffe JS, Cook EH, Jr., Geschwind D, Roeder K, Devlin B, State MW (2011) Multiple recurrent de novo CNVs, including duplications of the 7q11.23 Williams syndrome region, are strongly associated with autism. Neuron 70: 863–85.

Shansky RM, Murphy AZ (2021) Considering sex as a biological variable will require a global shift in science culture. Nat Neurosci 24: 457–464.

Song JY, Ichtchenko K, Sudhof TC, Brose N (1999) Neuroligin 1 is a postsynaptic cell-adhesion molecule of excitatory synapses. Proc Natl Acad Sci U S A 96: 1100–5.

Tabuchi K, Blundell J, Etherton MR, Hammer RE, Liu X, Powell CM, Sudhof TC (2007) A neuroligin-3 mutation implicated in autism increases inhibitory synaptic transmission in mice. Science 318: 71–6.

Toddes C, Lefevre EM, Brandner DD, Zugschwert L, Rothwell PE (2021) mu-Opioid Receptor (Oprm1) Copy Number Influences Nucleus Accumbens Microcircuitry and Reciprocal Social Behaviors. J Neurosci 41: 7965–7977.

Uchigashima M, Cheung A, Futai K (2021) Neuroligin-3: A Circuit-Specific Synapse Organizer That Shapes Normal Function and Autism Spectrum Disorder-Associated Dysfunction. Front Mol Neurosci 14: 749164.

Varoqueaux F, Jamain S, Brose N (2004) Neuroligin 2 is exclusively localized to inhibitory synapses. Eur J Cell Biol 83: 449–56.

Werling DM, Geschwind DH (2013) Sex differences in autism spectrum disorders. Curr Opin Neurol 26: 146–53.

Yoshida T, Yamagata A, Imai A, Kim J, Izumi H, Nakashima S, Shiroshima T, Maeda A, Iwasawa-Okamoto S, Azechi K, Osaka F, Saitoh T, Maenaka K, Shimada T, Fukata Y, Fukata M, Matsumoto J, Nishijo H, Takao K, Tanaka S, Okabe S, Tabuchi K, Uemura T, Mishina M, Mori H, Fukai S (2021) Canonical versus non-canonical transsynaptic signaling of neuroligin 3 tunes development of sociality in mice. Nat Commun 12: 1848.

